# Greater local diversity under older species pools may arise from enhanced competitive equivalence

**DOI:** 10.1101/2020.04.20.052316

**Authors:** Devin R. Leopold, Tadashi Fukami

**Affiliations:** Department of Biology, Stanford University; Department of Botany and Plant Pathology, Oregon State University

**Keywords:** biodiversity, chronosequence, co-occurrence, functional diversity, ecosystem age, local-regional, microcosm, species pool, species richness

## Abstract

Local ecological communities tend to contain more species when they are located within a geologically older region, a pattern that has traditionally been attributed to the accumulation of species in the regional species pool. In this explanation, local species interactions are assumed to play a minor role in the formation of the regional species pool, which is instead thought to be driven by speciation and dispersal occurring across larger areas. Here, we provide evidence suggesting a more important role of local species interactions than generally assumed. In an experiment in which we assembled 320 local communities of root-associated fungi under 80 species pools, we varied the species richness of the experimental species pools and the mean age of the sites from which we collected the fungal species across a 4-myr soil chronosequence in Hawaii. We found that realized local species diversity in the assembled communities increased more extensively with increasing species-pool richness when species were from older sites. We also found that older species pools had lower functional and phylogenetic diversity, indicating that the evolution of greater competitive equivalence among potential colonists enabled higher local diversity under older species pools. Our results suggest that the tendency of older regions to have higher local richness arises not simply because older species pools are more speciose, but also because the constituent species have evolved traits that allow them to co-occur more readily in local communities.

## Introduction

Processes occurring across large geographical scales, such as vicariance and allopatric speciation, can dictate local patterns of species diversity (1–4). For example, geologically older regions and historically more abundant habitats tend to be associated with greater local diversity (5–7). This relationship between the age of a region and the level of local diversity has traditionally been attributed to the accumulation of regional species richness over time (6, 8). However, in addition to having greater richness, species pools in older regions may also include taxa that have evolved traits that facilitate local coexistence, enhancing local diversity. Such traits may evolve as a result of coevolution among species in the same regional species pool (9–14).

If competitive interactions favored co-occurrence of species with greater trait differences in the past, niche filling through diversification and immigration could increase functional diversity in the regional species pool (8, 15–17). Greater functional diversity in the species pool could then contribute to greater local diversity by reducing trait overlap among members of the species pool. Alternatively, if the outcomes of local species interactions were primarily determined by a competitive trait hierarchy, convergence on traits that equalize species’ intrinsic fitness differences could ensue (18, 19). This convergence could also promote local diversity because reduced fitness differences would allow locally rare species to persist (20). These hypotheses that invoke traits that govern how species locally interact remain largely untested.

Determining whether the composition of older species pools contributes to greater local diversity requires comparisons that account for differences in species pool richness and other potentially confounding factors, such as differences in regional species abundance distributions (21), local extinction and colonization dynamics (22), and the rate of local disturbance (23). Plant-associated fungal symbionts, which are phylogenetically and functionally diverse and can be manipulated in controlled microcosm experiments, represent an underutilized yet tractable system to test hypotheses linking regional species pools to local community assembly.

In this study, we test the hypothesis that the accumulation of fungal symbiont diversity within a host plant will saturate with increasing species pool richness, but that greater local diversity will be realized when the region contributing to the species pool is older. To this end, we assembled local communities of root-associated fungi using experimental species pools that varied in species richness and the geological age of the sites contributing to each species pool. For this experiment, we used fungal isolates from the roots of an ericaceous plant, *Vaccinium calycinum*, collected from five sites across a 4.1-million-year soil chronosequence in Hawaii. In addition, we quantified functional and phylogenetic relationships among species to test the hypothesis that species pools from older ecosystems would have greater functional diversity, which should facilitate local species coexistence.

## Results

Fungal communities within *V. calycinum* seedlings accumulated diversity at a decreasing rate with increasing species pool richness (*χ*^2^_(1)_) = 10.0, *P* = 0.002; Fig. 1; Table S1). As hypothesized, the relationship between pool richness and realized local diversity varied with species pool age, with older pools showing increasingly higher diversity as regional species richness increased *t*_1,70_ = 2.60, *P* = 0.011; Fig. 1; Table S1).

**Figure 1:**
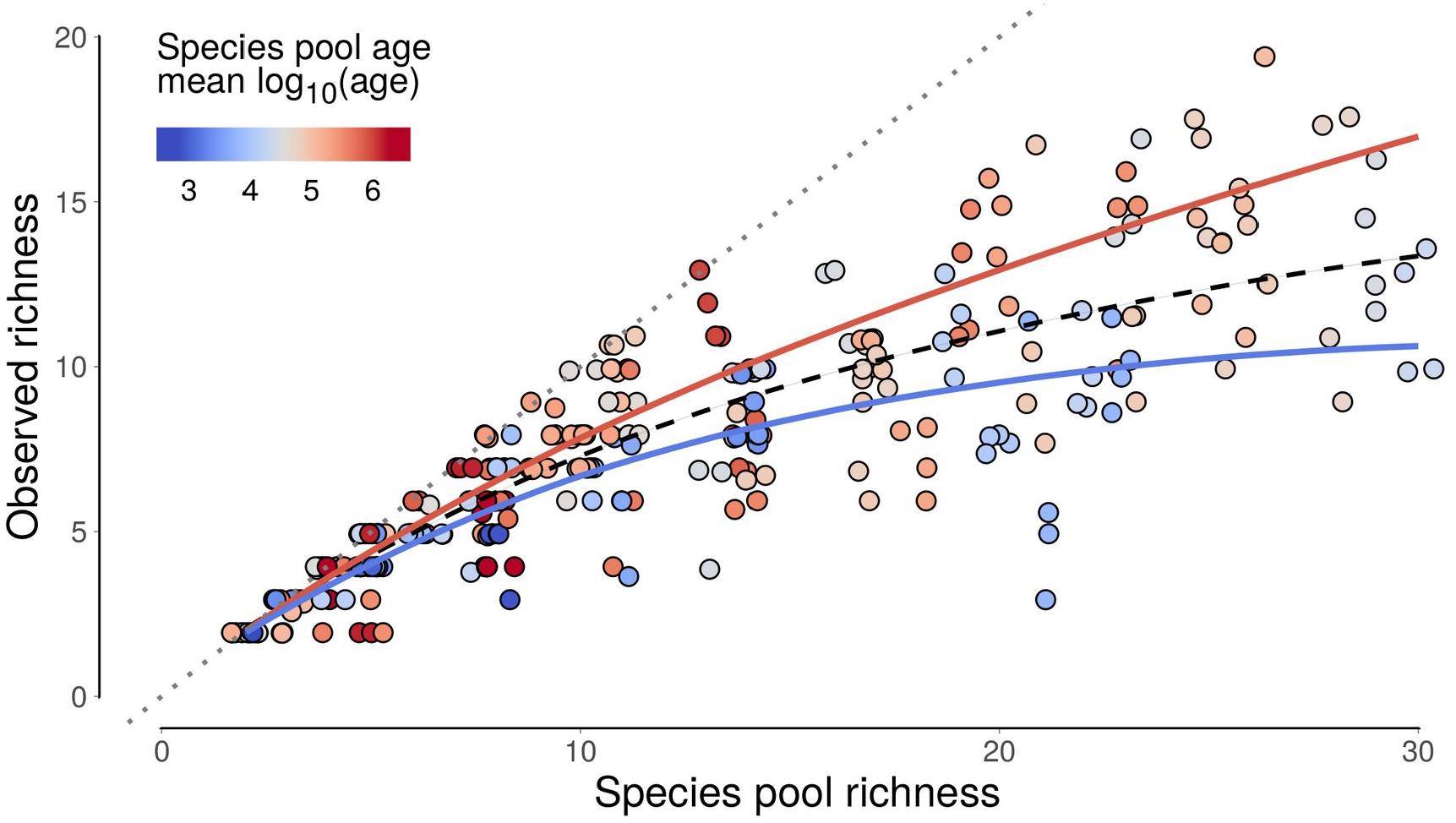
The relationship between species pool richness and local diversity (observed richness) in experimental microcosms, showing the theoretical maximum local diversity (dotted gray line) and a non-linear saturating model (a Michaelis-Menten function) fit to the data (dashed black line). Point colors indicate the mean of the log_10_-transformed age of the chronosequence sites represented in each species pool. Colored lines show the predicted effect of ecosystem age on the accumulation of diversity with increasing species pool richness.

We also found that the functional and phylogenetic diversity of the species pools decreased with increasing pool age (*t*_1,74_ = −4.92, *P* < 0.001; Fig. 2a; Table S2). Moreover, communities assembled with species pools from younger sites tended to become phylogenetically more dispersed during community assembly than communities associated with species pools from older sites, which tended to become phylogenetically more clustered (*t*_1,70_ = −4.24, *P* = 0.001; Fig. 3).

**Figure 2:**
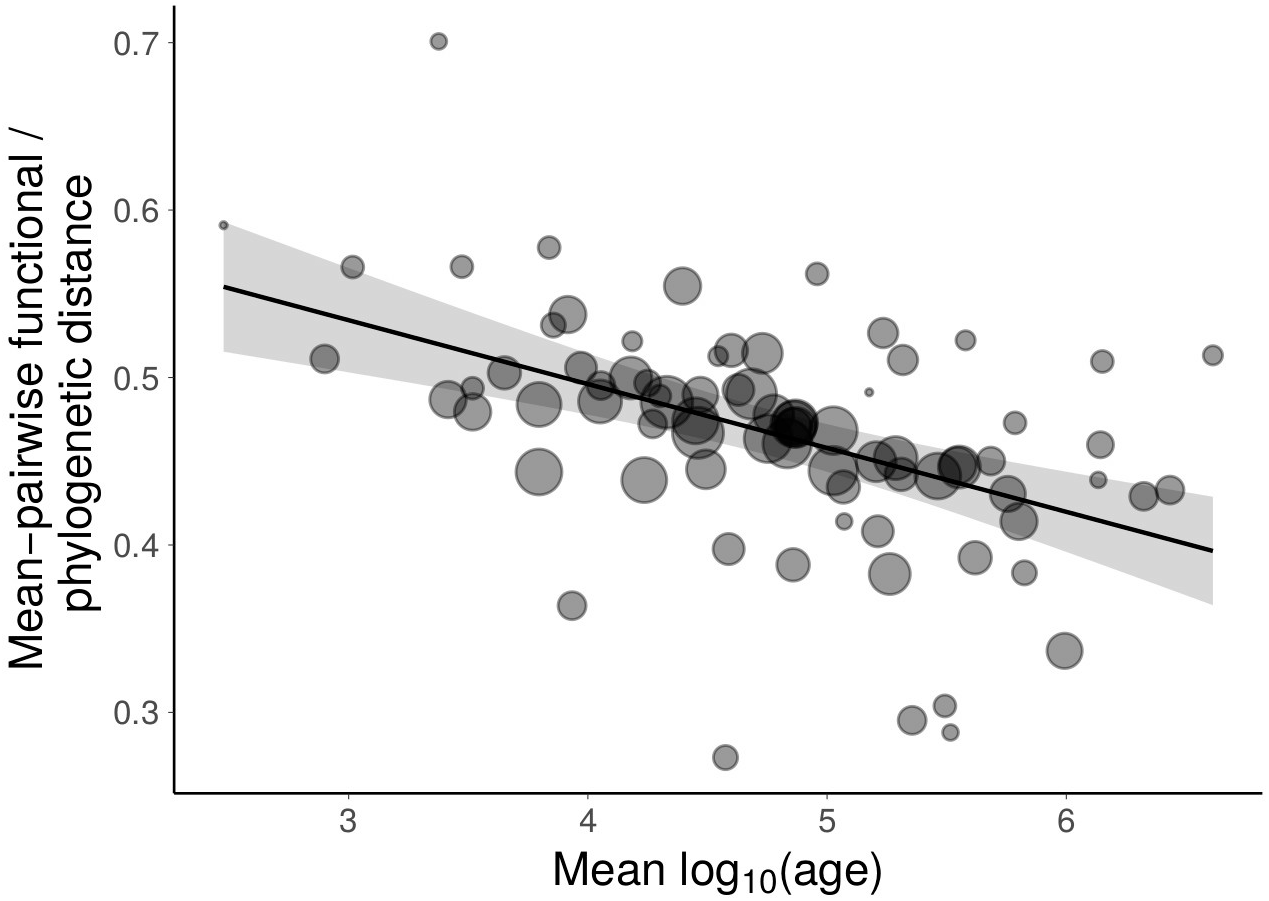
The relationship between the combined functional and phylogenetic diversity of the experimental species pools and the mean of the log_10_-transformed ages of the sites where the constituent species originated. Point size is proportional to species pool richness.

**Figure 3:**
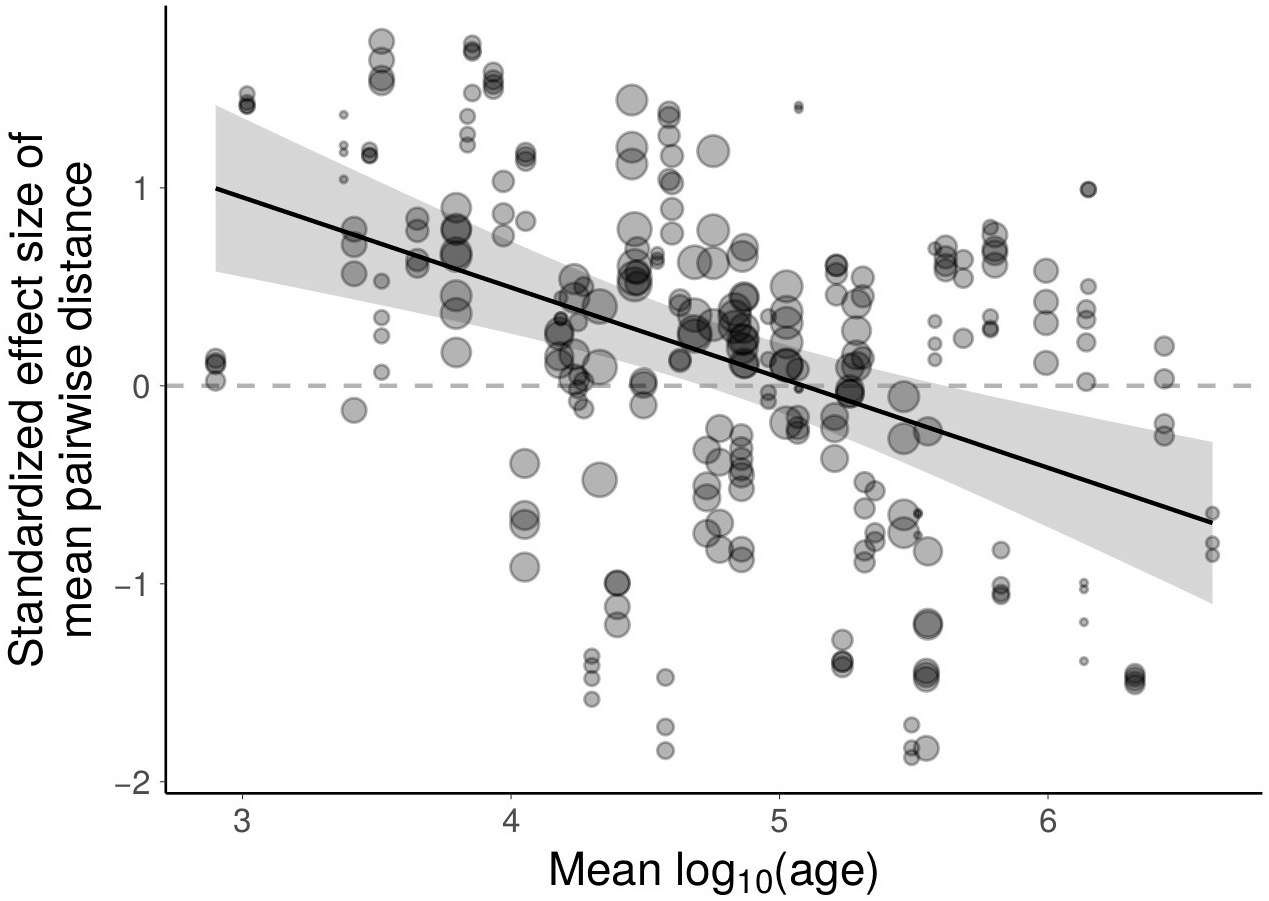
Effect of species pool age on the standardized effects size of mean pairwise functional and phylogenetic distance, showing that assembled communities had higher dispersion than the species pool if the pool was from young sites, whereas assembled communities had less dispersion than the species pool if the pool was from old sites. Point size is proportional to species pool richness.

## Discussion

Taken together, our results indicate that local diversity increases with the geological age of a region not just as a result of increased regional species richness, but also because the regional species pool contains species that are more likely to co-occur in local communities. Although the mechanisms responsible for fine-scale species co-occurrence in our microcosms are unknown, our results are consistent with evidence from molecular profiling of fungi in field-collected *V. calycinum* roots, which showed that the diversity and evenness of the fungal symbiont community associated with an individual plant increased with ecosystem age (24).

The finding that older species pools had lower functional and phylogenetic diversity (Fig. 2) suggests that increased coexistence of species from older sites may be the result of convergence on similar ecological strategies (i.e., equalizing fitness differences), rather than niche differentiation (18, 25). We do not know which traits determine the outcome of species interactions in this system, but both evolutionary and ecological mechanisms could contribute to trait convergence in older ecosystems. For example, a competitive trait hierarchy may lead to evolutionary convergence on similar root colonization strategies over geological time (19, 26, 27). It is also possible that changing conditions with long-term soil development led to trait convergence only in older sites. In plants, progressive phosphorus limitation with increasing soil age can result in a shift of species coexistence mechanisms from niche differentiation to trait convergence (28). Similar trends may exist in root-associated fungi, so that they coexist through niche differentiation in younger soils, but converge on a common resource-retentive strategy with declining phosphorus availability (29), resulting in coexistence through functional convergence and reduced fitness differences.

We cannot entirely rule out niche partitioning due to environmental heterogeneity as a mechanism facilitating diversity in the natural fungal communities we sampled for this experiment. For example, the accumulation of complex organic layers and recalcitrant nutrient pools during pedogenesis could promote diversity in ericaceous root-associated fungi through niche partitioning along nutrient resource axes (30). This fine-grained resource heterogeneity in older soils may facilitate coexistence among species in the regional species pool more than the broad scale environmental heterogeneity present across the chronosequence (31). However, soil nutrient resource partitioning is unlikely to explain the patterns we observed in the experimental microcosms, where a single growth media was used and nutrients were supplied in mineral form.

We did not detect a significant signal of clustering or dispersion in the phylogenetic or functional structure of the assembled communities. This lack of pattern could be due to the relatively small number of species used in our experiment and the biases introduced by our focus on ascomycetous fungi that could be easily cultured (32). In addition, spatial resource partitioning within roots between mycorrhizal fungi and non-mycorrhizal endophytes may not result in a strong phylogenetic signal due to the phylogenetic overlap among these functional groups (33, 34). Nonetheless, we did find that species pools from younger sites tended to increase in functional and phylogenetic dispersion during assembly more than species pools from old sites (Fig. 3). This result is consistent with a shift towards equalizing fitness differences, rather than niche differentiation, as the primary mechanism of species coexistence (28).

Greater local diversity in geologically older habitats has been thought to reflect increased richness of the regional species pool (6, 8). The novel finding of our study is that older species pools can also consist of species that are more likely to coexist. Our results are consistent with an increasing role of equalizing fitness differences in older species pools and a shift from niche partitioning in younger ecosystems to convergence on similar ecological strategies in older ecosystems, possibly in response to progressive phosphorus limitation. Increased local diversity as a result of strengthened species coexistence mechanisms might be a general feature of older species pools.

## Materials and methods

### Overview and experimental design

We isolated fungi from the roots of an ericaceous plant, *Vaccinium calycinum*, collected from 5 sites across a 4.1-million-year soil chronosequence in Hawaii, known as the Long Substrate Age Gradient (LSAG) (35). We then assembled experimental species pools of 2-30 fungi, manipulating geologic age by systematically varying the mean of the log_10_-transformed ages of the sites where individual species were collected. We also manipulated the variance of the log_10_-transformed site ages to account for the potential confounding effects of edaphic variation among sites on species pool functional diversity and assembly outcomes. We used site age variance as a proxy for edaphic variation because the LSAG chronosequence represents a progression of longterm soil development, resulting in greater differences in soil properties (e.g., physical structure, nutrient availability, etc.) with increasing differences in site age (35). A total of 80 experimental species pools (including one control treatment with no fungi) were inoculated onto sterile *V. calycinum* seedlings in individual microcosms, each replicated 4 times (320 total microcosms), and community assembly outcomes were assessed using Illumina metabarcoding after 5 months.

### Fungal culture collection

At each LSAG site, we sampled fungi from the roots of 12 randomly selected *V. calycinum* plants. A portion of the root system of each plant and the adhering soil (*ca*. 25 x 25 x 10 cm) was excavated with a hand trowel and transported immediately to the University of Hawaii, Hilo campus, where samples were refrigerated (4 °C) and processed within 24 hours. Fine terminal roots were manually separated from the soil and rinsed in tap water to remove all visible soil particles and 4 segments (*ca*. 2 cm) were haphazardly selected and pooled for each plant. Pooled root samples were surface-sterilized by sequential vortexing for 1 minute in sterile water, 70% EtOH, and 50% household bleach (4.5% available chlorine), followed by 3 rinses in sterile water. From each surface-sterilized root fragment, 2 segments (*ca*. 2 mm) were excised and aseptically transferred to separate petri dishes containing modified Melin-Norkans (MMN) agar, amended with the antibiotics gentamycin (15 mg l^-1^), streptomycin (15 mg l^-1^), and tetracycline hydrochloride (12 mg l^-1^) to inhibit bacterial growth. For each pair of excised root segments, one was placed on media that also contained benomyl (4 mg l^-1^), a fungicide that suppresses many ascomycetous fungi, facilitating the isolation of basidiomycetous fungi from roots (36, 37). Plates were monitored regularly, and hyphae growing from root segments were immediately transferred to new MMN plates and maintained by serial transfer every 4 to 8 weeks, depending on growth rate.

### Identifying fungal isolates

One representative of each fungal morphotype from each site was selected for possible use in the microcosm study and identified using molecular methods. Fungal DNA was extracted by scraping hyphae from the surface of a colonized agar plate and lysing cells with Extract-N-Amp Plant (Sigma-Aldrich) extraction and neutralization solutions. The complete nrDNA ITS gene and partial large subunit (28S) gene were PCR amplified using the primer pairs ITS1f-ITS4 (38, 39) and LROR-LR5 (40). PCR products were sequenced using single-pass Sanger sequencing (Beckman Coulter Genomics, Danvers, MA, USA) and sequences were deposited in GenBank under accession numbers MT321740–MT321793. Putative taxonomy was assigned by manually searching the full ITS sequence and the complete ITS and partial LSU sequences against the UNITE species hypothesis database v8.2 (41, 42).

We tentatively identified the fungal isolates used in the microcosm experiment as belonging to a range of ascomycetous taxa commonly associated with ericaceous plants, predominantly in the order Helotiales (Fig. S1). Many of the closest matching sequences in the UNITE database originated from the roots of other ericaceous species and were taxonomically affiliated with ericoid mycorrhizal fungi, including *Hyaloscypha bicolor, Oidiodendron maius, Pezoloma ericae* and poorly classified species within the Helotiales. Other taxa had affinities to groups of common endophytic fungi associated ericoid roots, including *Chaetosphaeria, Cladophialophora Phialocephala*, and *Rhizoderma veluwensis*, along with numerous unclassified Hyloscyphaceae. There is considerable taxonomic overlap between these two functional groups and the interactions between our isolates and the host plants are not well-documented (33, 34). However, given the aggressive surface sterilization protocol used and the taxonomic affiliations, we expect all taxa to be either ericoid mycorrhizal fungi or non-mycorrhizal endophytes with intimate associated with *Vaccinium calycinum*.

### Functional /phylogenetic characterization of fungi

Two complementary approaches were used to characterize the functional similarity among fungal isolates, direct measurements of functional traits and estimation of phylogenetic relationships. For functional traits, we quantified the relative abilities of each isolate to grow on 95 different carbon substrates using phenotypic microarrays (Biolog, FF microplate). Hyphal suspensions were prepared for inoculating microarrays by blending hyphae scraped from the surface of an agar plate in 60 ml sterile water. The hyphal suspension was then centrifuged at 3500 rpm for 5 minutes, decanted and the hyphae were resuspended in a sterile aqueous solution of 0.25% Phytagel and 0.03% Tween 40 to achieve a total volume of *ca*. 50 ml and an optical density of 0.06 at 590 nm. Using a multichannel pipette, 100 μl of this solution was transferred to each well of three replicate FF microplates for each fungal isolate. Plates were sealed with Parafilm to prevent desiccation and incubated at room temperature for 6 weeks. Fungal growth was monitored by reading optical density at 740 nm using a microplate reader (Spectramax 190, Molecular Devices), and measurements were collected at 1, 3 and 6 days, and weekly thereafter. Substrate use profiles for each species were generated by fitting a loess smoother to the optical density values in each well at each time point and integrating the area under the curve as a cumulative measure of substrate use. Values for the no-substrate control well were subtracted from all other wells to account for growth not attributable to the available carbon substrates.

To estimate phylogenetic relationships (Fig. S1), we use the previously sequenced 740-bp region of the nrDNA large ribosomal subunit (28S), which allows better resolution of phylogenetic relationships in ascomycetous fungi than the ITS region (43). We also retrieved a sequence of the matching gene region of *Saccharomyces cerevisiae* from GenBank [accession # KY109285.1] to use as an outgroup for phylogenetic analyses. Sequences were aligned on the T-Coffee webserver (44) and alignment columns were weighted using transitive consistency scoring (45) before phylogenetic tree construction using maximum likelihood and non-parametric, approximate likelihood-ratio tests of branch support with PhyML (46).

### Fungal species pools

Experimental species pools were constructed using a subset of the total culture library that excluded replicate isolates from the same site with identical nrDNA sequences. Initial Sanger sequencing revealed only 4 Basidiomycete taxa, which we excluded from the assembly experiment to avoid excessive influence of phylogenetic outliers in our analyses. Preliminary tests during methods development also revealed 4 pathogenic isolates (all *Cryptosporiopsis* spp.) that rapidly killed seedlings *in vitro;* these isolates were also excluded from the experiment.

In total, 54 fungal isolates were used in the construction of the experimental species pools. To ensure that individual isolates could be identified in the mixed communities at the end of the experiment, isolates with > 97% similarity in their ITS2 nrDNA gene sequences (the targeted barcoding sequence, see below) were never included in the same species pool. The compositions of species pools were selected using a stratified random sampling of the possible parameter space defined by the species pool richness (2-30 species) and the range of possible values of mean log_10_-transformed site age and variance of log_10_-transformed site age (age-variance), given the available species (Fig. S2). Briefly, 20 million random possible pools were generated and the parameter space was divided into 79 equal area bins, from which one species pool was randomly selected. This approach assured that more extreme parameter values would be sampled and allowed us to disentangle the effects of pool age from age-variance.

### Microcosms

Microcosms were assembled in 2.5 cm x 15 cm cylindrical glass culture tubes. Each tube received 3 g (dry) of a mixture of sphagnum peat, fine vermiculite and perlite (1:5:1) and 7 ml of a low-carbon mineral nutrient solution, following (47), containing 0.600 mM NH_4_NO_3_, 0.599 mM (NH_4_)2HPO_4_, 0.662 mM KH_2_PO_4_, 0.170 mM MgSO_4_.7H_2_O, 0.300 mM NaH_2_PO_4_, 0.080 mM K_2_SO_4_, 0.034 mM FeNaEDTA, 0.006 mM ZnSO_4_.7H_2_O, 3.69 μM MnSO_4_.H_2_O, 0.04 μM Na_2_MoO_4_.2H_2_O, 9.55μM H_3_BO_3_, 1 μM CuSO_4_.5H_2_O and 100 mg l^-1^ glucose. Culture tubes were fitted with polypropylene closures, modified to include 3 2-mm-diameter vent holes which were covered with 2 layers of micro-pore tape to allow gas exchange. Assembled microcosms were autoclaved twice for 30 minutes at 121 °C with a 24 hour interval and allowed to cool in a sterile-airflow hood.

### Seedlings

*V. calycinum* fruits were collected at the 300-year LSAG site and were cleaned, dried and refrigerated before use (48). Prior to assembling the microcosms, seeds were surface-sterilized by gentle vortexing in a 10% solution of H_2_O_2_ for 5 minutes, rinsing in sterile water, and then germinated in petri plates on 0.8% water agar in a lighted growth chamber. Individual 6-week-old seedlings (*ca*. 1 cm tall) were aseptically transferred into the sterile microcosms, which were sealed with Parafilm. Microcosms were then placed in a lighted growth chamber providing 150 μmol m^-1^ s^-1^ of photosynthetically active radiation with lighting cycle of 16 h light (25 °C) and 8 h dark (20 °C). The positions of the microcosms within the growth chamber were randomized once a week throughout the experiment and seedlings were allowed to acclimate after being transferred to the microcosms for 3 weeks before fungi were added.

### Fungal inoculation

Fungal inocula consisted of hyphal suspensions prepared in individual 1-pint canning jars, each fitted with a stainless steel blender assembly, filled with 80 ml distilled water and autoclaved for 30 minutes at 121 °C. Once cool, each sterile jar received 4 agar plugs (1 cm) taken from the growing edge of a 4 week old fungal culture, and was blended at high speed for 60 seconds. Inoculum for each species pool was prepared in a sterile 15 ml centrifuge tune by combining 150 μl of the hyphal suspension of each individual isolate in the pool and adding sterile water for a total of 5 ml. Microcosms were inoculated under a sterile air-flow hood and received 1 ml of a mixed-species inoculum (or sterile water) which was applied at the base of the seedling.

### Microcosm harvest

Microcosms were harvested 5 months after the introduction of the fungal species pools. Roots were separated from shoots and were gently cleaned of growth media in molecular biology grade water using flame sterilized forceps and briefly dried on sterile filter paper. The fresh weight of roots and shoots were recorded. Shoots were dried at 60 °C and analyzed for C and N concentration using an elemental analyzer (Carlo Erba NA 1500). After weighing, roots were transferred to sterile 2-ml screw cap tubes and stored at −80 °C. Empty 2-ml tubes (4) were opened while harvesting roots and carried through all stages of the metabarcoding procedure as a negative control. We did not find any evidence for an effect of species pool composition on seedling biomass or leaf chemistry and present a graphical summary of these results as supplemental material (Fig. S3).

### Fungal species composition

Fungal species composition within seedlings at the end of the experiment was determined using Illumina metabarcoding, targeting the ITS2 region of the nrDNA gene. This region was identified by Sanger sequencing as being variable enough to discriminate between isolates while having minimal length variation to minimize amplicon sequencing bias (49). In addition to the microcosm samples, we included samples that consisted of DNA from pure cultures of each fungal isolate. Including samples with DNA from individual isolates allowed verification of the Sanger sequencing of the target barcoding region.

DNA was extracted from the entire root system of each seedling using bead beating and the Nucleomag 96 Plant Kit (Macherey-Nagel) on a KingFisher Flex (ThermoFisher) automated magnetic particle processor. A dual indexed, two-stage fusion-PCR procedure was used for Illumina library preparation, following the primer design of (50). We modified the stage-one primers to include the gene primers 58A1F (51) and ITS4. Because these primers amplify both fungal and host ITS2 nrDNA, we designed an amplification-blocking oligo (52) that overlapped with 2-bp at the 3’ end of the forward primer, extended into a host specific sequence, and terminated with a 3’ C3 spacer to block elongation during PCR (5’-GTA GCG AAA TGC GAT ACT TGG-3SpC3-3’). Mismatches between the blocking oligo and fungal DNA were consistent across all species, limiting the probability of introducing amplification bias. Stage-one PCR reactions (25 μl) were carried out using MyTaq Hot-Start Red Mix (Bioline), 10 μM forward and reverse primers, 100 μM blocking oligo, 1 μl template DNA, and a thermocycler program consisting of an initial cycle of 95 °C (3 min), 32 cycles of 95 °C (20 sec), 60 °C (10 sec), and 72 °C (10 sec), followed by a final elongation cycle at 72 °C (5 min). Stage-two PCR reactions (25 μl) used 2 μl of the stage-one PCR product as template, 10 μM primers, and a thermocycler program consisting of an initial cycle of 95 °C (3 min), 6 cycles of 95 °C (20 sec), 55 °C (10 sec), and 72 °C (10 sec), followed by a final elongation cycle at 72 °C (5 min). Final PCR products were cleaned and normalized to 2.5 ng μl^-1^ using Just-a-Plate, 96-well normalization and purification plates (Charm Biotech), then pooled and sent to the Stanford Functional Genomics Facility for 250-bp paired-end sequencing on the Illumina MiSeq platform. The resulting raw data waspublicly archived in the NCBI Sequence Read Archive (BioProject PRJNA613615).

### Bioinformatics

Bioinformatic processing of the Illumina sequence data involved first removing gene primers and amplicon length heterogeneity spacers with cutadapt v1.18 (53). Initial quality assessment indicated poor quality for the reverse reads which were not used in subsequent analyses. The forward reads were cropped to 200 bp, filtered at a maximum expected error rate threshold of 2 bp, and denoised with the R-package DADA2 (54) to identify the unique amplicon sequence variants present in each sample. Priors, consisting of the expected amplicons for each sample (i.e. only isolates present in the experimental species pool), were provided to facilitate accurate denoising. Because the initial denoising did not fully resolve all single species control samples to a single cluster, amplicon sequence variants within each sample were further collapsed using the R-package stringdist (55) and a Hamming distance threshold of 4 bp. Unexpected reads in microcosm samples were predominantly host DNA or likely contaminants from the sample processing and library preparation procedure, which were present in very low abundance and often also identified in the no-DNA negative control samples. These reads were filtered from the data before analysis. Read counts for each denoised amplicon in each sample were merged into a species-by-sample matrix for analyses, discarding samples with low sequencing depth (< 5000 reads). After bioinformatic processing of the metabarcoding data, 286 samples remained, leaving 3 or 4 replicates of 75 unique species pools.

All bioinformatics code has been publicly archived (DOI: 10.5281/zenodo.3758735).

### Statistical Analyses

To quantify local diversity we used multiple rarefaction of the community sequencing data, sampling without replacement to the minimum sequencing depth across all samples (5736 sequences), and calculating the mean species richness over 1000 iterations. To determine if the relationship between local diversity and species pool richness was linear or curvilinear (i.e., saturating), we used linear mixed-models in the R-package nlme (56), including species pool composition as a random effect. To account for heteroscedasticity we allowed the model variance to increase with increasing species pool richness. Support for a curvilinear model was tested by comparing models with and without a polynomial term using a likelihood-ratio test.

We used two approaches to test whether the relationship between species pool richness and local diversity was affected by species pool age. First, we used polynomial mixed-models, as above, testing for a significant interaction between species pool richness and age. To account for a possible effect of site heterogeneity we also included age-variance and the interaction between age-variance and species pool richness. In the second approach, we fit a generalized non-linear least squares model with nlme, using a Michaelis-Mentin function, which takes the form,

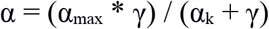

where α is local diversity and γ is species pool richness. The estimated parameters, α_max_ and α_k_, represent the asymptotic value of α and the value of γ at which half of the asymptotic value of α is reached, respectively. As with the polynomial mixed-model approach, we allowed the model variance to increase with species pool richness. We then modeled the standardized (Pearson) residuals of the non-linear model as a function of species pool richness and age to test whether the residuals varied systematically with species pool age. Specifically, we fit a linear mixed-model with species pool composition as a random effect and interactions between species pool richness and both species pool age and age-variance as fixed effects. We then used likelihood ratio tests to simplify the model, removing both the interaction between pool richness and age-variance (*χ*^2^_(1)_ = 0.061 *p* = 0.81) and the direct effect of age-variance (*χ*^2^_(1)_ = 0.16 *p* = 0.68). We then refit the final model using restricted maximum likelihood estimation to test the significance of the remaining fixed-effects. Conclusions from the polynomial mixed-model and analysis of the generalized non-linear model residuals were identical. We present the results of the generalized non-linear model here because the functional form more closely approximates the expected saturating relationship between regional richness and local diversity.

To test the relationship between phylogenetic or functional diversity and species pool composition, we first quantified the mean pairwise phylogenetic and functional distances among isolates for each species pool. Phylogenetic distances were square-root transformed to account for non-linear scaling between evolutionary and functional distances (57). Carbon substrate use data was reduced to the first 10 principle components with PCA (accounting for ~75% of total variation) to collapse highly correlated substrates. Euclidean distance of the PCA coordinates was then used to calculate mean pairwise functional distances among isolates for each species pool. Because patterns were broadly similar for phylogenetic and functional diversity, we combined these metrics with equal weight, following the method of (58), resulting in a single measure of functional and phylogenetic diversity, FPdist. We used multiple linear regression to test the hypothesis that mean pairwise FPdist increases with species pool age, accounting for both species pool richness and site heterogeneity (age-variance).

To test whether changes in phylogenetic and functional diversity during community assembly (i.e. increased clustering or dispersion) were related to species pool age we calculated the standardized effect sizes for the mean pairwise FPdist, using the R-package picante (59). We weighted species abundances using Hellinger-transformed community sequencing data and used a null model where taxa names were permuted 999 times on the distance matrix. The standardized effect sizes were modeled as a response using a linear mixed-effects model, with species pool richness, age, and age-variance as fixed effects and species pool composition as a random effect.

All R code and the associated sample meta data required to recreate the analyses presented here has been publicly archived (DOI: 10.5281/zenodo.3758735).

## Acknowledgments

Laboratory space and other resources used in this research were generously provided Michael Shintaku and Anne Veillet, at the University of Hawaii, Hilo, Scott Gibb, at the USDA-ARS, Pacific Basin Agricultural Research Center, and Christian Giardina, at the USFS, Institute of Pacific Islands Forestry. This research was funded by a National Science Foundation Doctoral Dissertation Improvement Grant (grant no.1600521).

