## Supplemental materials for "Greater local diversity under older species pools may arise from enhanced competitive equivalence"

**Table S1:** Analysis of the relationship between species pool richness and local diversity, showing (a) the estimated parameters and 95% confidence intervals for the generalized non-linear least squares model (Michaelis-Menten) and (b) the effects of species pool richness, age, and their interaction on the standardized residuals from the Michaelis-Menten model.

**a) Parameters of Michaelis-Menten model**

| Parameter | Estimate | LCI | UCI |
|---|---|---|---|
| $\alpha_{max}$ (asymptote) | 22.9 | 19.9 | 25.9 |
| $\alpha_k$ (half-max) | 21.3 | 17.0 | 25.6 |

**b) Analysis of model residuals**

| Parameter | Estimate | SE | DF | *t* | *p* |
|---|---|---|---|---|---|
| Richness | -0.126 | 0.049 | 74 | -2.561 | 0.013 |
| Age | -0.025 | 0.034 | 74 | -0.712 | 0.479 |
| Richness:Age | 0.028 | 0.011 | 74 | 2.605 | 0.011 |

**Table S2:** Results of multiple-linear regression testing whether the species pool mean-pairwise combined functional and phylogenetic distance varied with species pool age, richness and site heterogeneity (age-variance).

| **Parameter** | **Estimate** | **SE** | ***t*-value** | ***p*-value** | **partial-$r^2$** |
|---|---|---|---|---|---|
| Intercept | 0.625 | 0.040 | 15.396 | < 0.001 | |
| Richness | -0.001 | 0.001 | -0.861 | 0.392 | 0.01 |
| Age-variance | 0.014 | 0.005 | 2.997 | 0.004 | 0.11 |
| Age | -0.037 | 0.008 | -4.923 | < 0.001 | 0.25 |

**Table S3:** Results of linear mixed-model testing whether the degree of functional (and phylogenetic) dispersion or clustering during community assembly changed with species pool age, accounting for species pool richness and age-variance.

| Parameter | Estimate | SE | DF | *t*-value | *p*-value |
|---|---|---|---|---|---|
| Intercept | 2.057 | 0.572 | 199 | 3.585 | < 0.001 |
| Richness | 0.003 | 0.012 | 70 | 0.298 | 0.767 |
| Age-variance | 0.077 | 0.063 | 70 | 1.152 | 0.253 |
| Age | 0.440 | 0.105 | 70 | -4.248 | 0.001 |

**Figure S1:** Estimated phylogenetic relationships of the 54 fungal isolates. The maximum likelihood tree was constructed with PhyML, using the first 740 bp of the nrDNA large ribosomal subunit. Colors indicate the age of the chronosequence site where each isolate was collected. For each isolate, the OTU id is followed by the GenBank accession number of of the associated sequence, and then the putative taxonomic assignment, based on comparison of the full nrDNA internal transcribe spacer region against the UNITE v8.2 database.

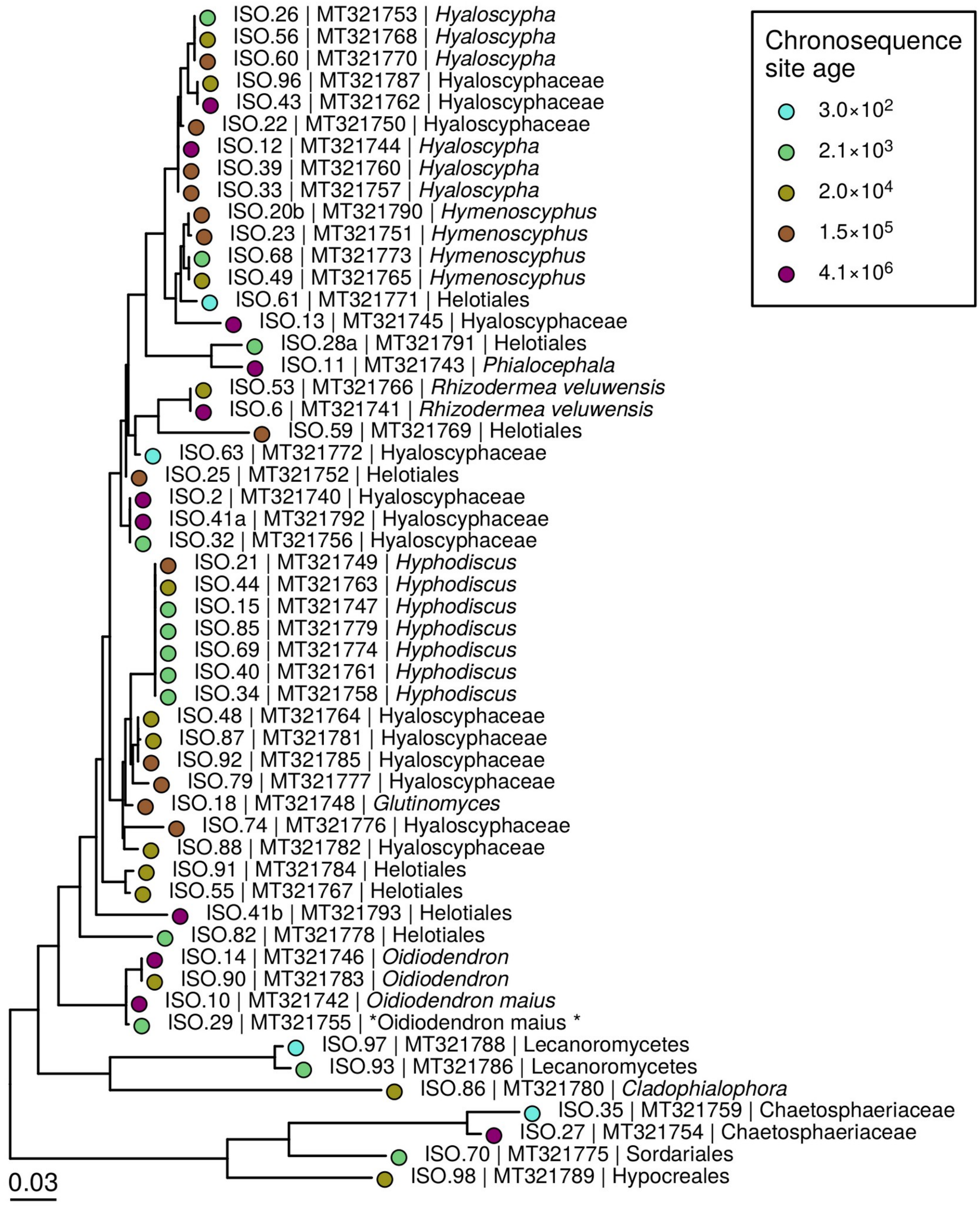

**Figure S2:** Graphical summary of experimental pool design, showing the pairwise relationships between species pool richness (a & b), and the mean (a & c) and variance (b & c) of the $\log_{10}$-transformed ages of the sites included. Light gray points indicate the parameter values for 20 million randomly generated possible species pools and black points show the values for the 79 species pools used in the experiment.

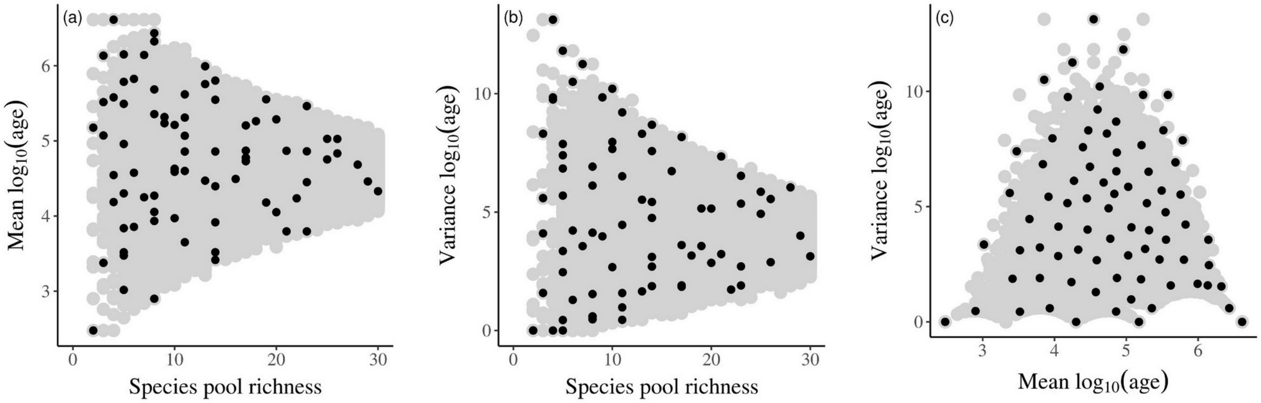

**Figure S3:** Relationships between experimental manipulations of species pool composition and seedling (a-c) shoot and (d-f) root biomass and leaf tissue (g-i) nitrogen and (j-l) carbon concentrations. Horizontal lines indicate the median values for the four control seedling that were grown without fungi.

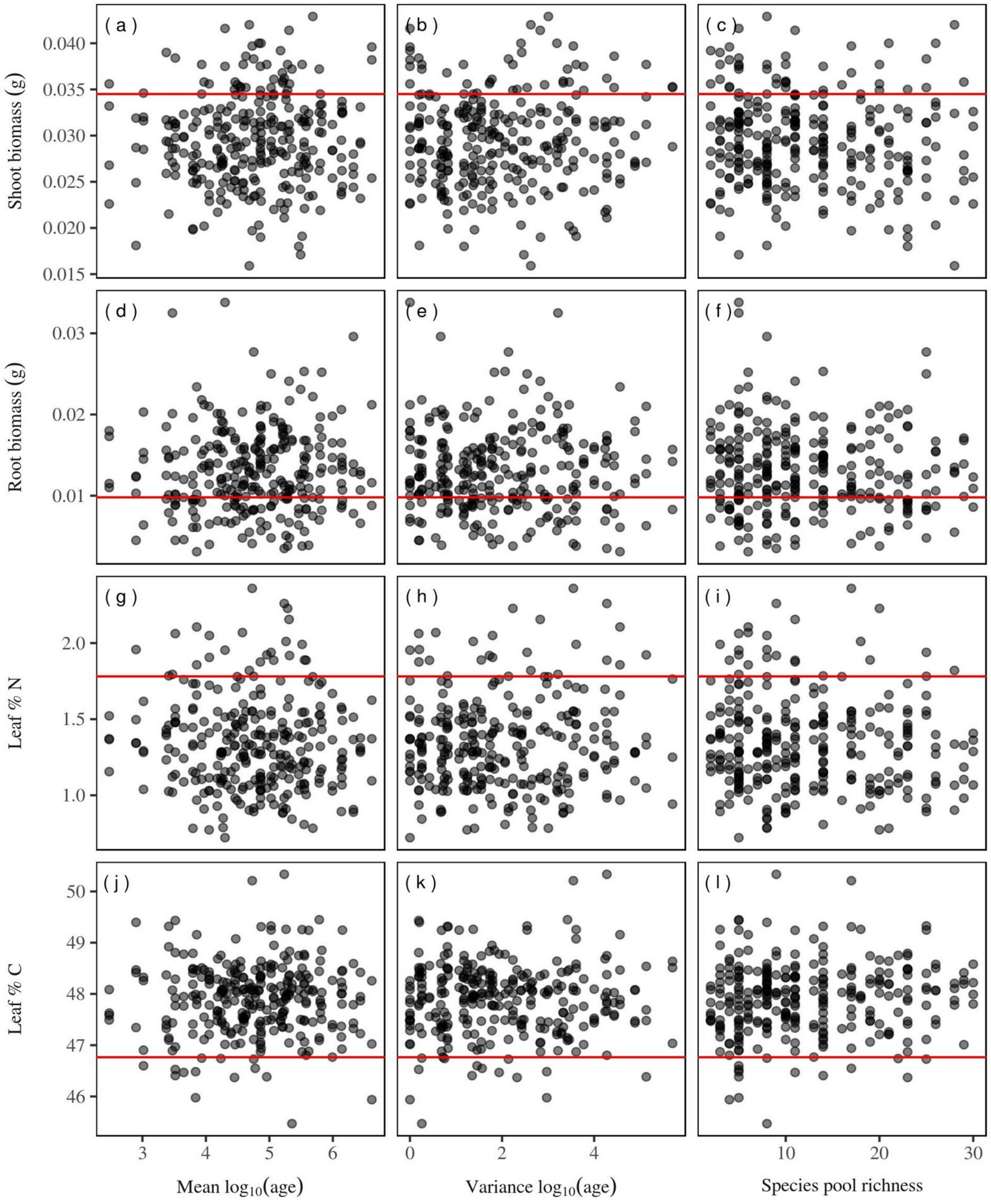
